# Multichromatic Near-Infrared Imaging to Assess Interstitial Lymphatic and Venous Uptake *In Vivo*

**DOI:** 10.1101/2021.03.07.434298

**Authors:** Fabrice C. Bernard, Jarred Kaiser, Sarvgna K. Raval, Zhanna V. Nepiyushchikh, Thanh N. Doan, Nick J. Willett, J. Brandon Dixon

**Affiliations:** Wallace H. Coulter Department of Biomedical Engineering, Georgia Institute of Technology and Emory University, Atlanta, GA; Department of Orthopaedics, Emory University, Atlanta, GA; George W. Woodruff School of Mechanical Engineering, Georgia Institute of Technology, Atlanta, GA; Department of Orthopaedics, Atlanta Veteran’s Affairs Medical Center, Atlanta, GA; Parker H. Petit Institute for Bioengineering and Bioscience, Georgia Institute of Technology, Atlanta, GA

**Keywords:** Tissue Optics, NIR Imaging, Venous, Lymphatic, Clearance

## Abstract

**Significance:** Changes in interstitial fluid clearance are implicated in many diseases. Using NIR imaging with properly sized tracers could enhance our understanding of how venous and lymphatic drainage are involved in disease progression or enhance drug delivery strategies.

**Aim:** We investigated multichromatic NIR imaging with multiple tracers to assess *in vivo* microvascular clearance kinetics and pathways in different tissue spaces.

**Approach:** We used a chemically inert IR Dye 800CW (free dye) to target venous capillaries and a purified conjugate of IR Dye 680RD with a 40 kDa PEG (PEG) to target lymphatic capillaries *in vivo*. Optical imaging settings were validated and tuned *in vitro* using tissue phantoms. We investigated multichromatic NIR imaging’s utility in two *in vivo* tissue beds – the mouse tail and rat knee joint. We then tested the ability of the approach to detect interstitial fluid perturbations due to exercise.

**Results:** In an *in vitro* simulated tissue environment, free dye and PEG mixture allowed for simultaneous detection without interference. Co-injected NIR tracers cleared from the interstitial space via distinct routes allowed assessment lymphatic and venous uptake in the mouse tail. We determined that exercise after injection transiently increased lymphatic drainage as measured by lower normalized intensity immediately after exercise, while exercise pre-injection exhibited a transient delay in clearance from the joint

**Conclusions:** NIR imaging enables of simultaneous imaging of lymphatic and venous-mediated fluid clearance with great sensitivity and can be used to measure transient changes in clearance rates and pathways.

## Introduction

The circulatory system maintains tissue homeostasis through the continuous delivery of nutrients and oxygen to the tissue space and the removal of proteins and waste products. Crucial to this process is the removal of interstitial fluid, proteins, and lipids by the lymphatic vasculature; this fluid then returns to the circulation through absorption at the lymph nodes and delivery to the central venous system through the lymphatic ducts. Recent developments in optical imaging have provided new capabilities to quantify lymphatic function *in vivo*.

Generally, there are two routes of fluid clearance from tissues: 1) venous uptake and 2) lymphatic uptake. Venous return in tissue beds is passive, size-dependent, and varies based on capillary physiology^1,2^. In contrast, lymphatic capillaries originate from the tissues and have flap-like openings, which non-discriminately allow molecules of all sizes to enter. The extrinsic motion of the surrounding tissue combined with the intrinsic contractility of downstream lymphatics, create transient pressure gradients that allow fluid and macromolecules to enter the vessel and be transported downstream. Impaired interstitial fluid clearance has been implicated in various diseases, including lymphedema^3^, cancer^4^, and arthritis^5^. Techniques to measure clearance kinetics from interstitial spaces are critical to evaluating disease state and different tissues’ ability to drain molecules from the interstitial spaces. These measurements have been assessed classically via radiolabeled agents, which carry potential toxicity and require additional safety measures ^6–8^. However, the advent of near-infrared (NIR) fluorescent imaging allows for cost-effective, high resolution, clinical and preclinical imaging in a variety of applications^9–11^.

NIR-based technologies have advanced considerably in the last decade—both in terms of imaging components as well as tracers and fluorophore-based probes—which have allowed for significant new *in vivo* capabilities. The NIR imaging window includes the visible and infrared light spectrum from 650-1300 nm, which due to longer wavelengths, penetrates tissues deeper than higher energy light^12–14^. Contrast agents like indocyanine green (ICG), polyethylene glycol (PEG) conjugated with NIR dyes, or NIR quantum dots have been used to visualize lymphatics and blood vessels *in vivo* ^15–19^. Preclinical NIR imaging has also previously enabled the measurement of tracers’ differential uptake as a function of size from different tissue beds^17,20^. Multichromatic NIR imaging (e.g., imaging with multiple NIR fluorescent probes) empowers mapping of the drainage zones of lymph nodes in rodents^19,21,22^. However, this has not yet been widely extended to differentiate between venous uptake and lymphatic uptake simultaneously in the same tissue bed.

We have previously shown the size-dependent uptake of 2 and 40 kDa NIR PEG into the venous and lymphatic circulation, respectively, after injection into the rat knee joint^23^. In that study, we demonstrated that the intra-articular injection of endothelin-1 (ET-1), a vasoactive compound in lymphatics and veins, transiently reduced the outflow of both PEG tracers from the joint in a dose-dependent manner. Due to these experiments’ monochromatic nature, we were unable to assess lymphatic and venous drainage simultaneously. The inability to differentiate clearance mechanisms and function between the venous and lymphatic systems is a critical technological gap that has broad implications for many different tissues and disease states. Coupling *in vivo* delivery with multichromatic NIR imaging could allow for the advancement of the understanding of how the venous and lymphatic drainage may change in the context of diseases or physical interventions. The objective of this manuscript was to develop a novel technological approach that couples NIR imaging with the size-dependent clearance of tracers *in vivo*. We hypothesized that a multichromatic imaging approach for differentially imaging the lymphatic and venous systems would show the technique’s utility in both the mouse tail, where the superficial vessels can be clearly visualized, and in the rat knee joint, where uptake occurs slowly and in deeper tissue structures. Additionally, we perturbed the joint microenvironment by exercising the rats on a treadmill and detected changes in venous and lymphatic clearance within the knee joint.

## Methods

### Tracers for *in vivo* injection

IR Dye 800CW carboxylate (free dye) (LI-COR Biosciences) was purchased as a dry lyophilized powder. 20 nanomoles were resuspended in 100 µl of sterile saline to make a 20 mM stock solution and used within a few days of resuspension. For tail injections, 2.5 µl of the stock solution was used. For tissue phantom studies and knee injections, the stock solution was diluted to 0.4 mM in sterile saline.

40 kDa methoxy polyethylene glycol (PEG) amine (JenKem Technology) was purchased as a dry lyophilized powder for PEG tracer synthesis. To conjugate 40 kDa PEG amine to IR Dye 680RD, 16 mg of PEG amine was reacted with 30 µl of 10 mg/ml IR Dye 680RD NHS ester (diluted in dimethyl sulfoxide (DMSO)) in a total of 1 ml of Dulbecco’s Phosphate-Buffered Saline (DPBS) overnight. Unreacted IR Dye, salts, and DMSO were removed via centrifugal filtration using deionized water and a 10 kDa molecular weight cutoff centrifugal filters (Amicon Ultra). After centrifugation, the purified tracers were separated into 10 equal volumes of 100 µl and aliquoted to 1.6 mg of tracer per aliquot, lyophilized, and kept frozen at -20°C for long term storage. For tail injections, aliquots were resuspended in 100 µl of sterile saline, and 2.5 µl of P40D680 was injected intradermally. For tissue phantom studies and knee clearance studies, aliquots were diluted to 1 mg/ml.

### Optical properties of tracers

To quantify each tracer’s absorbance in the visible and NIR range, 0.4 mM, and 1 mg/mL of the free dye and 1mg/mL of PEG were loaded into a standard UV/vis spectrophotometer (Ultraspec 2100, Biochrom). The emission and excitation spectra of free dye and PEG were assessed using a microplate reader with filter-based emission and detection capabilities (Synergy H4, BioTek). For PEG and free dye, fixed emission filters of 720 and 840 nm were used while sweeping the excitation source from 400 – 700 and 400 – 820 nm, respectively, to generate excitation curves. Following the excitation sweeps, fixed excitation wavelengths of 660 and 760 were used to conduct an emission sweep from 680/780 – 900 nm for PEG and free dye.

### NIR imaging setup

Multichromatic NIR imaging was carried out using a customized NIR setup ^17,24^. Briefly, the system consists of a cooled EMCCD camera (Evolve eXcelon, Photometrics) attached to a stereomicroscope with adjustable zoom (MVX10, Olympus), a shutter-controlled xenon arc light source (Lambda LS, Sutter Instrument Company), and a manual filter wheel equipped with standard Cy5.5 and ICG-B filter cubes (Chroma Technology). The electronic shutter was left open during continuous imaging sessions, and images were acquired using MicroManager software ^25^.

### Dye and tracer characterization and tissue phantom studies

Polydimethylsiloxane (PDMS) tissue phantoms, measuring 2 and 4 mm in thickness, were created using a previously described protocol^15^. By weight, 88.10% silicone elastomer base (Sylgard 184, Dow Corning) was mixed with 8.81% curing agent (Sylgard 184, Dow Corning), 1.76% Aluminum Oxide (Sigma Aldrich), and 1.32% cosmetic powder (Max Factor Crème Puff Deep Beige 42). PDMS phantoms were poured into plastic molds and left to cure in the oven at 60°C overnight.

To obtain reference measurements for our NIR imagine setup, stock PEG and free dye were diluted serially in two-fold dilutions in PBS. In separate 1.5 ml centrifuge tubes, free dye and PEG were diluted using PBS to 0.4 mM and 1 mg/ml. The tissue phantoms were used to demonstrate the effect of tissue depth on tracer intensity with the previous serially diluted samples. To simulate increasing tissue depth each centrifuge tube was imaged with no tissue phantom or with a 2- or 4-mm tissue phantom. To quantify the sensitivity to each tracer in the presence of the other NIR tracer, each tracer was diluted using the stock solution of the other tracer. The 2 mm tissue phantom was used to mimic the typical depth of the superficial collecting lymphatics in rodents. All images were taken with an exposure time of 50 and 5 milliseconds, respectively, for free dye and PEG.

### Tail injections to visualize and quantify routes of tracer clearance

To visualize the tail lymphatics and blood vessels, 20 µl of 1% (w/v) Evans blue solution was injected into the tip of the tail of an anesthetized mouse. Evans blue binds to interstitial proteins and is mainly taken up by lymphatics when injected intradermally. Post-euthanasia the skin was removed at the base of the tail to reveal the underlying vasculature. Images of the vasculature were taken using a standard color camera to provide a comparison with NIR images. For NIR imaging through the skin, isoflurane was used to anesthetize C57Bl/6J mice, and the animal was placed in a recumbent position (on its side). A mixture containing 2.5 µl of the free dye and 2.5 µl of PEG was mixed and loaded into 1 mL insulin syringes (Becton Dickinson) and injected intradermally into the tip of the tail. After injection, standard tape was gently applied to the base and the end of the tail to minimize drift from motion artifact during imaging. Free dye and PEG signals were imaged in 2-minute increments by manually changing the filter wheel to select the appropriate filter set every 1200 frames (Figure 1a). The free dye and PEG signal were evaluated at 50 and 20 milliseconds, respectively, and images were captured at ten frames per second.

**Figure 1.**
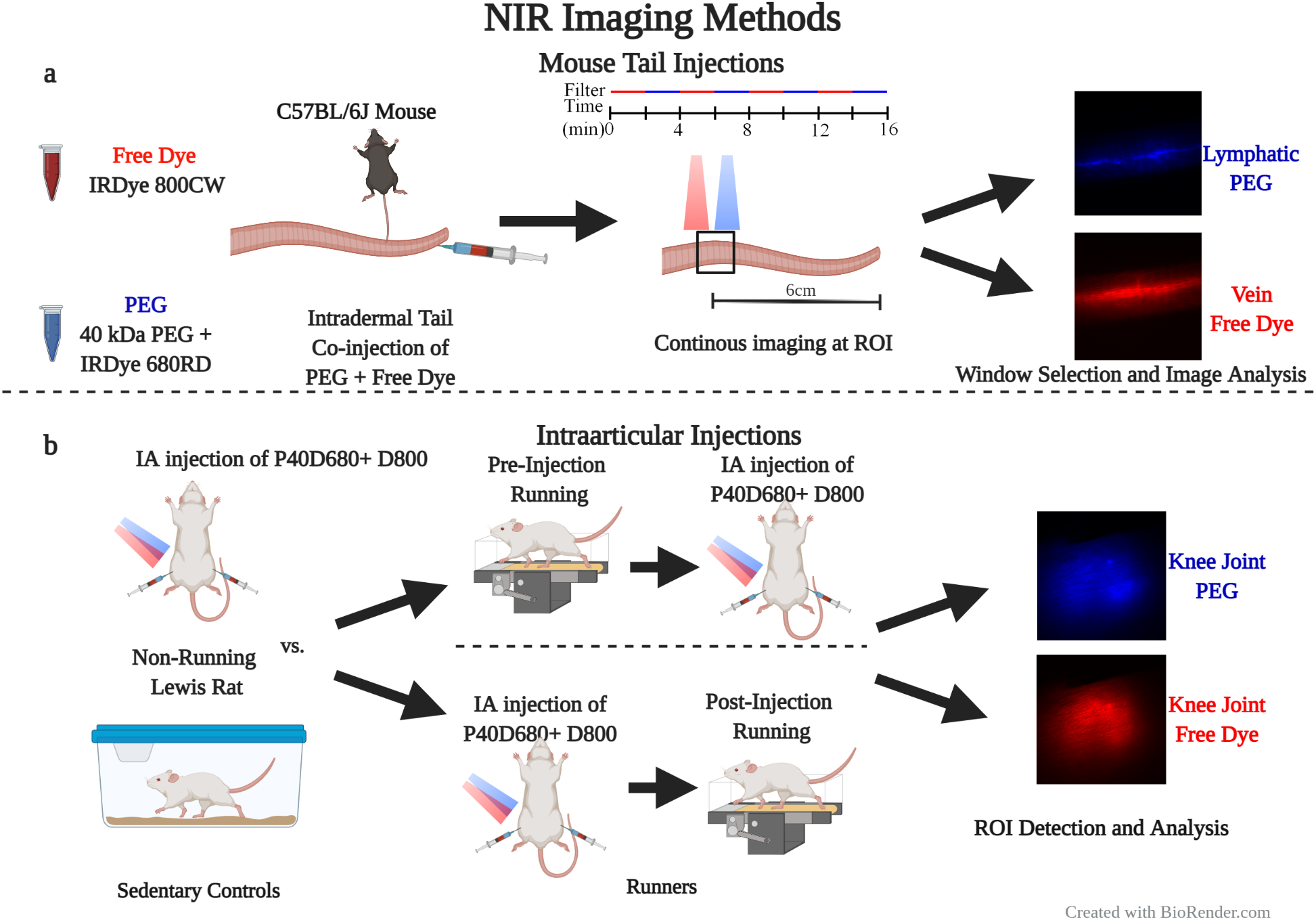
*In vivo* NIR imaging methods. **a**. Mouse tail injection and imaging methods. **b**. Intra-articular injections for assessing the effect of exercise.

Animal care and experiments were conducted under the institutional guidelines of the Georgia Institute of Technology. Experimental procedures were approved by the Georgia Institute of Technology Institutional Animal Care and Use Committee (IACUC).

### Intra-articular injections for clearance

Male Lewis rats weighing 350-400 grams were trained to run on the treadmill over two weeks. On day one, the rats were acclimated to the treadmill for 30 minutes without running. On day two, the treadmill was speed was set to 5 m/min for 5 minutes and 0 m/min for 25 minutes. Each day the time spent running was increased by 5 minutes a day until the rats could run for 30 minutes on consecutive days after two weeks. Rats that failed to walk the targeted duration twice over the training course were excluded from the study. All other rats were then randomly selected for the experimental procedure to either run or serve as controls for the study duration. Three sets of experiments were conducted with these two rat groups: 1) rats that did not run on the treadmill and were co-injected bilaterally with NIR tracers (No Running), 2) rats that were run on the treadmill for 30 minutes before injection (Pre-Injection Running), 3) rats that were run on the treadmill for 30 minutes immediately after injection (Post-Injection Running).

The day before the experiment, all rats were anesthetized, the hair was removed from the knees and lower abdomen, and background images of the knees were taken. Before imaging, each rat was induced via 5% isoflurane on the day of the experiment, which was maintained at 2% after induction. Tracers were injected in both knees and imaged at set time intervals (approximately 0, 1, 2, 3, 5, 7 12, & 24 hours) over the course of 24 hours.

Animal care and experiments were conducted per the institutional guidelines of the Atlanta Veteran Affairs Medical Center (VAMC). Experimental procedures were approved by the Atlanta VAMC IACUC.

### Image processing and analysis

Images captured using our custom NIR imaging system were saved as 16-bit depth 512 x 512-pixel TIF file format. For both tissue phantom experiments and *in vivo* knee joint clearance experiments, the tracer’s intensity in the image was quantified using a custom MATLAB (MathWorks) script. The ROI for each image was quantified by averaging the 5% highest pixel intensities. This ROI visually corresponded with the size and position of the knee space shown in Figure 1b. Data points were fitted to a monoexponential function f(t) = y0 + Ae^-kt^, where y0 is the offset, t is the time in hours, A is the normalized peak fluorescence at the maximum intensity, and k is the time constant. τ (tau) was determined as the inverse of the time constant. To compare each intervention’s short-term effects, we calculated the mean change in ROI intensity over the first hour and subtracted the mean value of internal control rats. To determine each intervention’s overall effects, we calculated the time constant for each runner and normalized it to the non-runner group’s mean.

For mouse tail injections of NIR dyes, the filter was changed manually every 2 minutes. To remove imaging artifacts from the manual changing of the filter, 60 seconds (600 frames) of each imaging window were cropped. Fiji software was used to crop, register, and quantify regions of interest (ROIs)^26^. An ROI was drawn on the blood and lymphatic vessels to monitor the signal of free dye and PEG over the experiment.

### Data presentation and statistics

All clearance data are presented as mean ± SEM. A Brown-Forsythe test was used to quantify if variances were significantly different. A student’s t-test with a Welch’s correction was used to compare venous and lymphatic area under the curve and tau in control rats. A one-way ANOVA with Dunnett’s multiple comparison test was used to calculate statistical significance for exercise studies.

## Results

### Optimization and characterization of NIR tracers using an *in vitro* tissue phantom

Absorbance spectra of free dye and PEG display an absorption maximum of 765 nm and 672 nm, respectively (Figure 2a). Emission and excitation spectra for these tracers also show each tracer’s expected maxima referenced to the full-width half maximum of the filter sets on the imaging system (Figure 2b). To determine the limits of detection in our imaging system as a function of concentration and tissue depth, we used tissue phantoms. Individual tracers were imaged in 1.5 ml centrifuge tubes with and without 2- and 4-mm phantoms. Increasing the tissue phantom thickness decreased the fluorescent intensity, though even at 4 mm, the dyes could be detected at a concentration of 3% of the injection concentration (Figure 2c). At a thickness of 2mm, which is within the depth of most superficial lymphatics in rodents, this detection limit was less than 1% of the injection site’s intensity. Notably, mixing the tracers did not affect the sensitivity to detect one dye when contained in the background of the other tracer (Figure 2d).

**Figure 2.**
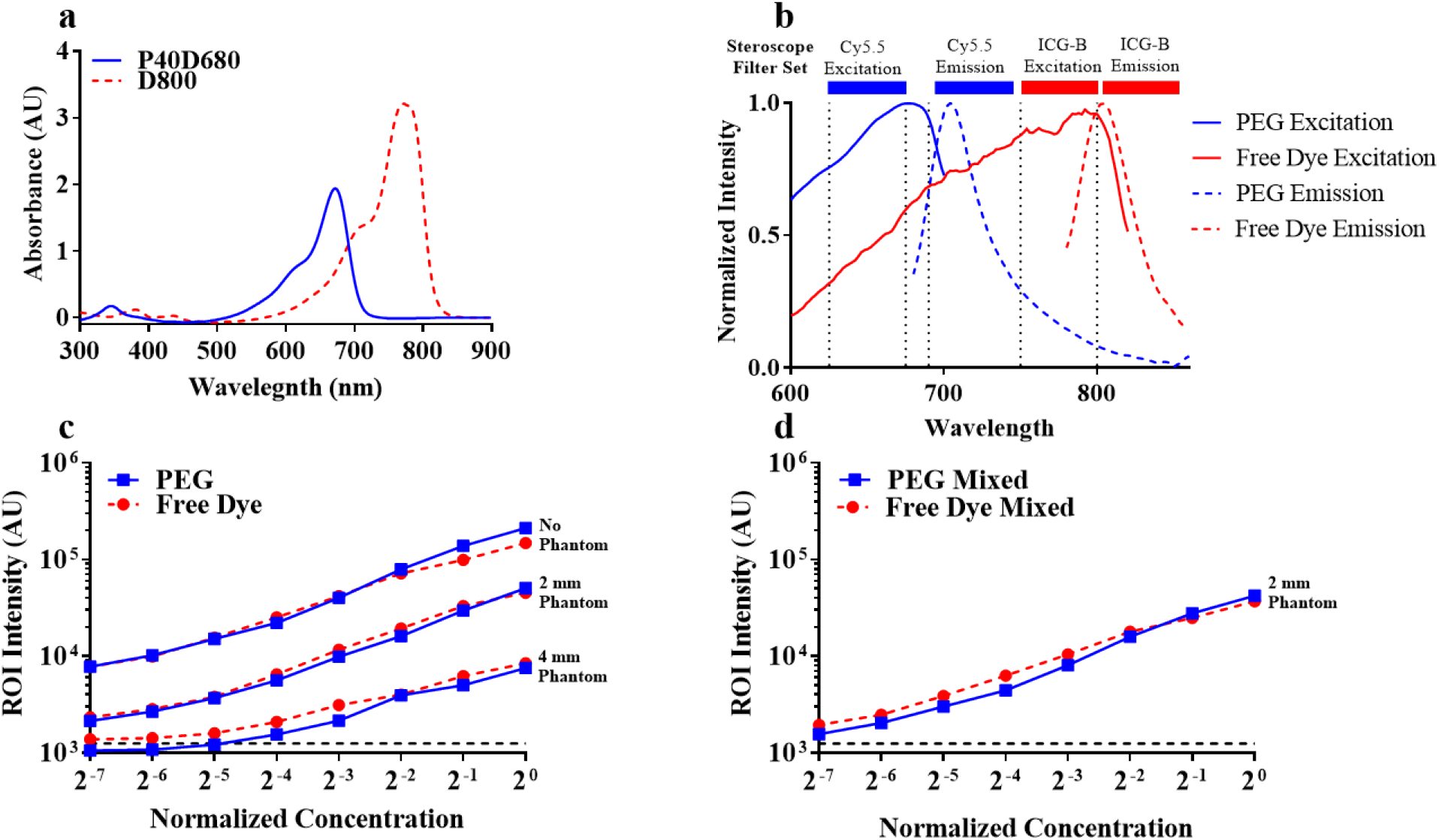
Sensitivity Analysis of Near-Infrared Dyes with Tissue Phantoms. **a**. Absorbance spectra show a unique absorption profile for each tracer. **b**. Solid and dashed lines show the excitation/emission spectra for 800CW carboxylate (free dye) and 680RD 40 kDa PEG (PEG), respectively. Our NIR stereoscope filter cube setup is represented by the bars above the graph, showing that our optical configuration is designed to read each tracer’s unique signal. **c**. Tissue phantoms used to determine the effect of tissue depth on detecting free dye and PEG showed reduced intensity as a function of phantom depth and serial two-fold dilution. Consequentially, we were able to see a reduction in signal intensity below the background (dotted line) and, therefore, could reach the limit for these tracers at this tracer at 25 ms exposure and 100 ms exposure time respectively, using a 4 mm tissue phantom. **d**. Free dye was serially diluted using a stock solution of PEG and vice versa. There was no change in overall intensity and sensitivity for each tracer due to mixing and imaging in a 2 mm phantom.

### NIR tracers of different size exit through spatially distinct clearance pathways

The mouse tail’s unique circulatory and lymphatic vasculature was visualized via intradermal injection of Evans blue and skin removal. The lymphatics immediately cleared Evans blue allowing clear visualization of the two lymphatic vessels running parallel to the tail vein and artery (Figure 3a). To simultaneously quantify lymphatic and venous drainage, these tracers were co-injected in the mouse tail. Images for each tracer were captured during two-minute imaging windows (eight recordings per tracer). Figure 3b shows that the routes of clearance of free dye in the blood circulation (red) and PEG dye in the lymphatics (blue) 10 minutes post-injection are spatially distinct and match the expected physiology where two lymphatic vessels flank a blood vessel. The free dye was initially detected in the lymphatic within the first imaging window (Supplementary Video 2) as the free dye is not expected to be excluded by lymphatics. The free dye intensity increased between the first, third, and fifth imaging windows and remains constant by the seventh (Figure 3d). However, PEG entered the lymphatics after injection and did not appear in the bloodstream over this time interval, demonstrating the lymphatic specificity of PEG (Supplementary Video 1). Also, the lymphatic tracer exhibited strong transient peaks in the signal intensity due to intrinsic phasic lymphatic contractions. In contrast, no such peaks were present in the tracer taken up into the blood circulation.

**Figure 3.**
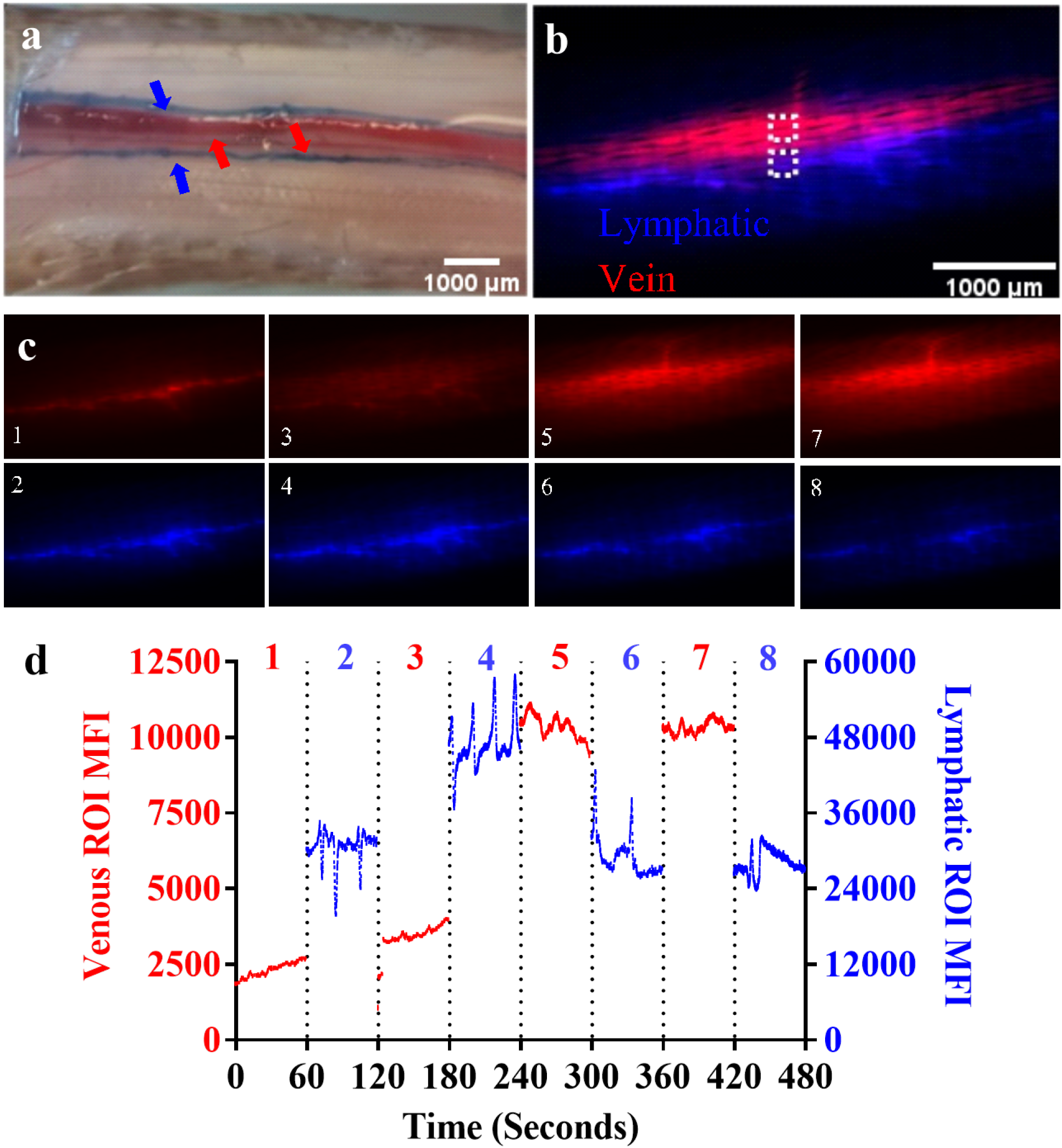
Co-injection of NIR tracers results in differential uptake of 800 CW Carboxylate and 680RD PEG 40 kDa. **a**. Evans blue dye injected into the tail of a mouse immediately before euthanasia shows the concentration of Evans blue dye in the lymphatics (blue arrows) that flank the blood vessels (red arrows) (scale bar = 1000 um). **b**. Free dye and PEG show the uptake of each NIR tracer in vein and lymphatics 10 minutes post injection (scale bar = 1000um). **c**. Free dye can initially be seen in the lymphatic; however, over time free dye concentrates in the circulation revealing the tail vein. PEG shows the sustained uptake of PEG dye into lymphatics. **d**. Measurement of signal intensity over the course of the four imaging windows for each respective tracer shows large phasic lymphatic contractions for the PEG tracer and increasing free dye signal over time.

### Co-injection to assess differential tracer clearance in the joint

The effect of exercise on intra-articular clearance has not been extensively studied^27,28^, specifically in quantifying the change in venous and lymphatic drainage. Therefore, after confirming size-dependent uptake from the tail, we used multichromatic imaging to assess intra-articular drainage. Unlike intradermal injections, materials from the joint space clear much slower ^29–31^; therefore, venous and lymphatic clearance from this interstitial depot was expected to occur over one day. Simultaneously injected tracer clearance PEG and free dye profiles (Figure 4a) exhibited an initial increase followed by monoexponential clearance kinetics consistent with previously reported figures^23^. Specifically, lymphatic tracers showed a more substantial increase in intensity after the injection, whereas this increase was less pronounced for the venous tracer. By 12 hours post-injection, the intensity of the free dye was not detectable (Figure 4a). The normalized area under the curve (AUC) for free dye was calculated to be 3.59 ± 0.18 and 18.57± 1.33 for PEG (p<0.0001), demonstrating significant retention of the PEG in the joint space (Figure 4b). The time constant (Tau) for the clearance of free dye was calculated to be 4.28 ± 0.25 hrs, while the time constant for PEG was 7.11 ± 0.51 hrs (p=0.0003) (Figure 4c). A detectable amount of the PEG tracer remained in the joint space even after 24 hours (Figure 4a), likely due to some PEG tracer remaining trapped in the interstitium.

**Figure 4:**
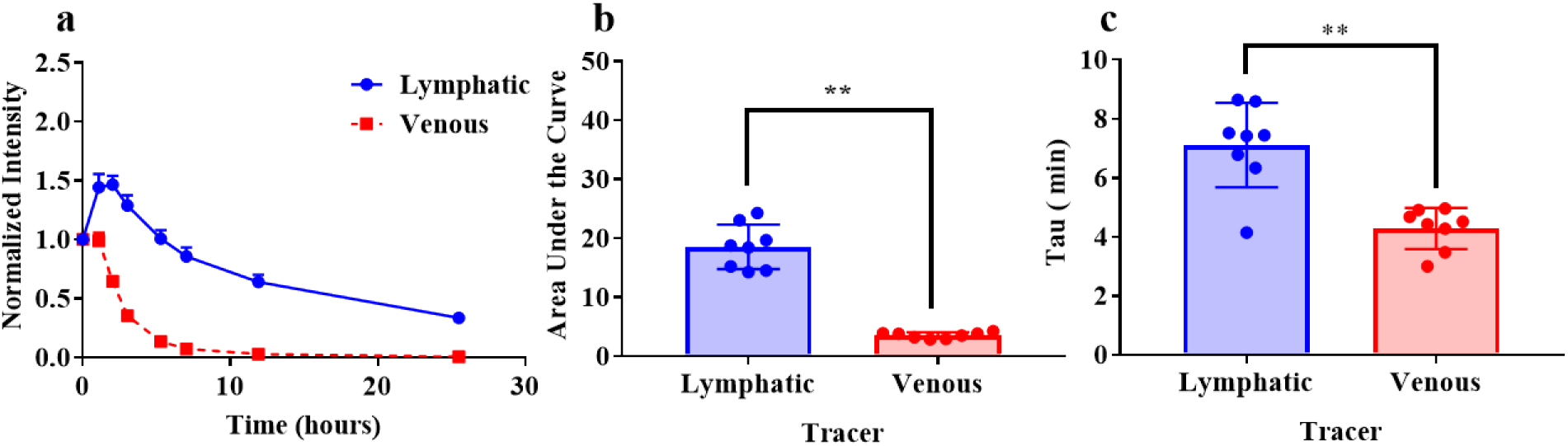
Co-injection of NIR tracers allows for the simultaneous detection of lymphatic and vascular mediated clearance from the joint space. **a**. Clearance profiles for PEG and free dye show the characteristic lymphatic and venous clearance, respectively. **b**. The areas under the curves show a significantly lower AUC for free dye compared to the PEG. **c**. First-order clearance constant tau was calculated for each tracer and is significantly higher for the lymphatic draining PEG vs the venous draining free dye.

### Multichromatic imaging for measuring patterns in joint clearance

To demonstrate the utility and sensitivity of multichromatic imaging to evaluate differential changes in clearance mechanisms for venous and lymphatic drainage, animals were exercised either pre- or post-injection. In rats that received no running intervention, clearance curves for both lymphatic and venous tracers had clearance profiles as expected. (Figure 5a). Qualitatively, from the initial injections, there did not appear to be differences in differences in the traces of animals that were designated runners and control animals. To quantitatively evaluate these differences, the normalized changes in fluorescence intensity were calculated (Fig 6a, c), as well as the time constant (Fig 6b, d).

**Figure 5.**
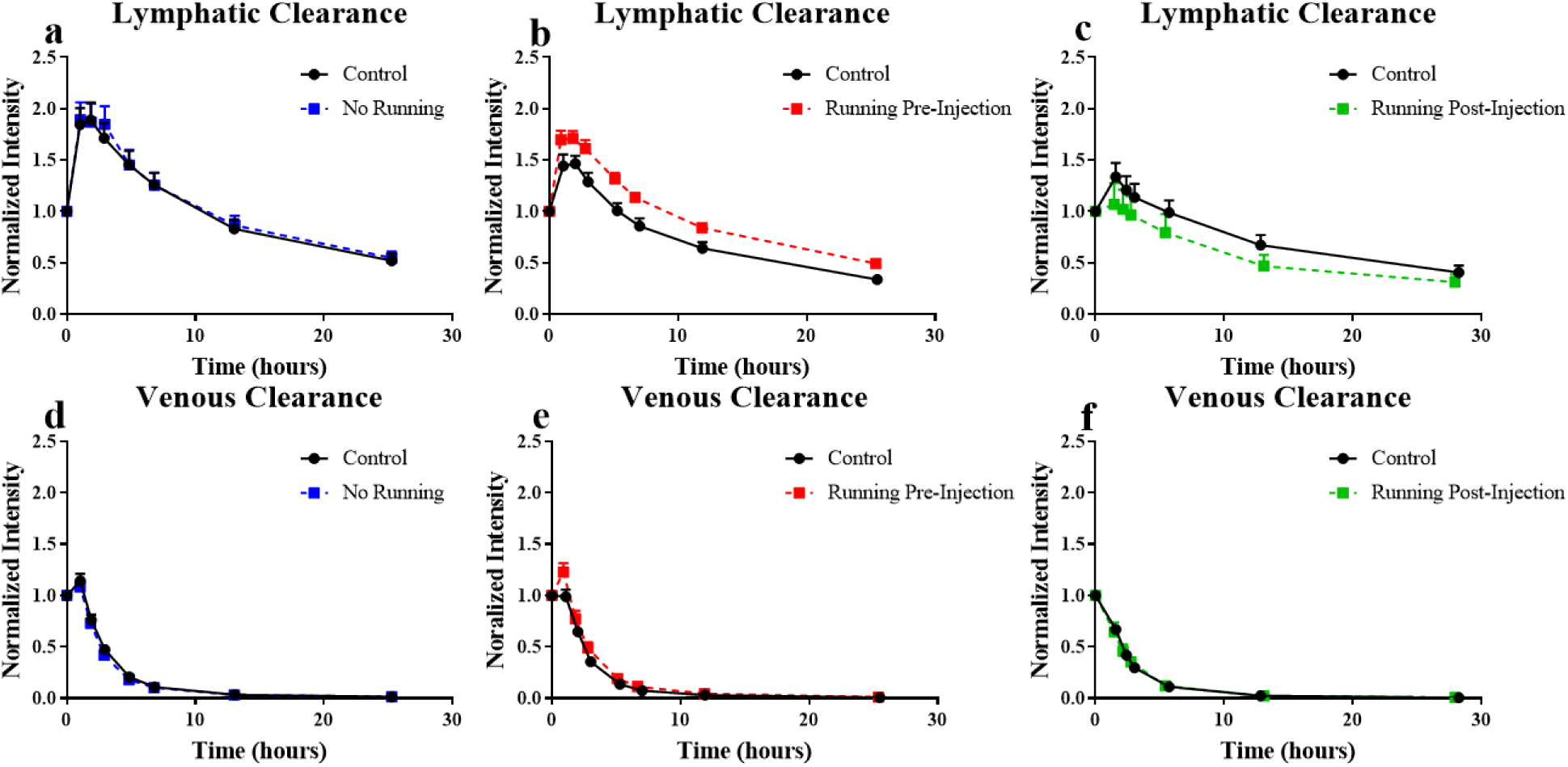
Clearance Profiles for PEG and free dye with running. **a-c**. Clearance profile of lymphatic specific tracer (PEG) for no running, pre-injection running and post-injection running experiments. **d-f**. Clearance profile for venous draining (free dye) for No running, pre-injection running, and post-injection running experiments.

**Figure 6.**
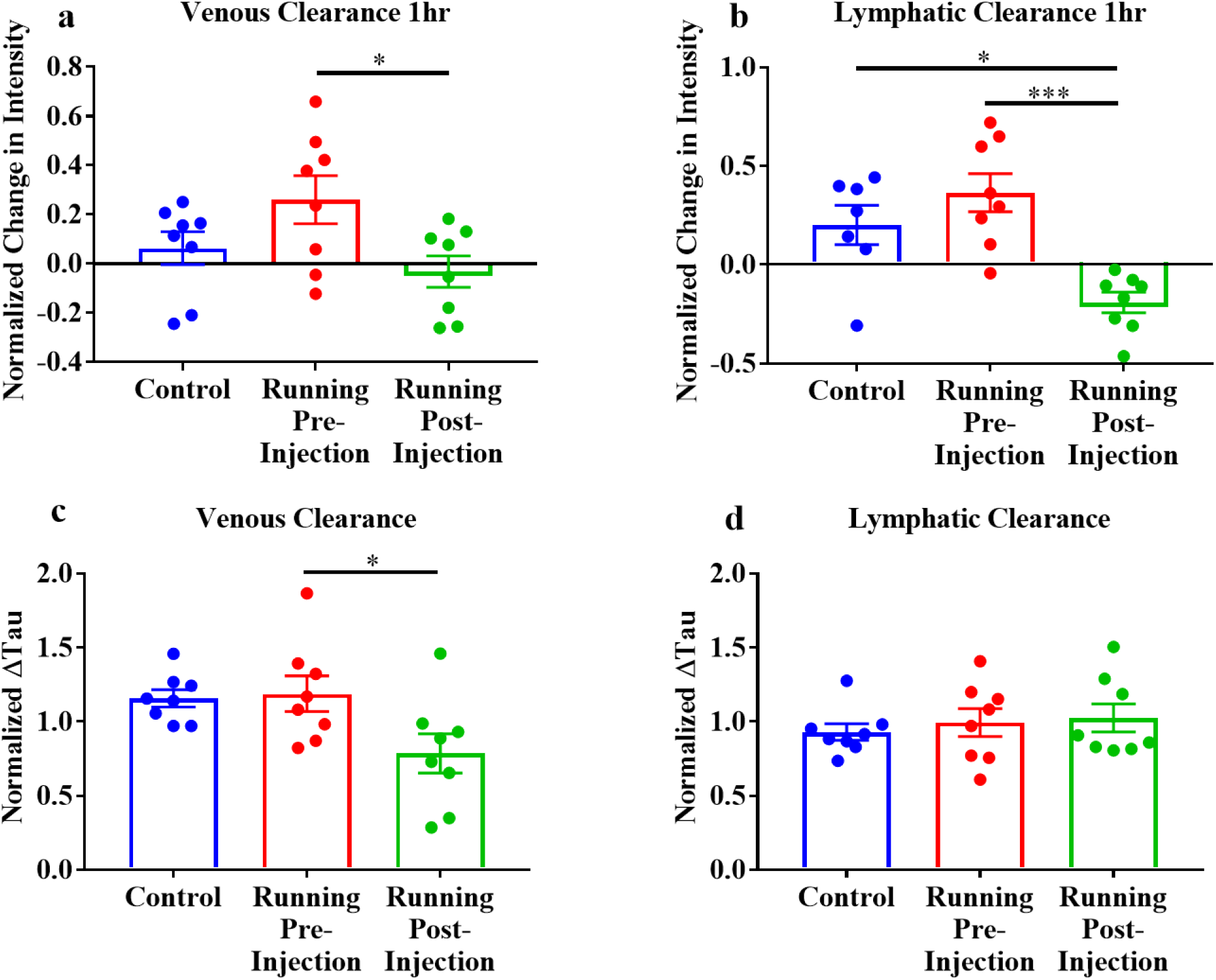
Effect of running on Normalized Change in Intensity and Normalized Tau **a, b**. The normalized intensity was calculated to assess the transient effect of running on dye dispersion in the joint. For each experiment, all runners were normalized to the mean of non-running controls. Compared to the control experiment running pre- or post-injection did not significantly change the initial free dye intensity; however, at the second captured timepoint, post-injection normalized intensity was significantly decreased compared to pre-injection running (*, p = 0.04). After 1 hour, lymphatic intensity was significantly decreased compared to controls (*, p = 0.01) and pre-injection running (***, p = 0.0003) **c, d**. Time constant (tau) for each condition was normalized to the controls for each day. Compared to intraexperimental non-running controls, there are no significant differences in clearance rate for either tracer. Compared to running pre-injection, clearance rate is significantly decreased (*, p = 0.04).

For venous drainage, running pre-injection (0.26 ± 0.10) significantly increased (p = 0.04) the change in fluorescence intensity compared to running post-injection (−0.03 ± 0.06), however neither were significantly different from control (0.06 ± 0.06) (Figure 6a). Figure 6c shows the normalized tau (venous) for running post-injection was calculated to be (0.79 ± 0.13) which is significantly reduced (p = 0.04) the compared to pre-injection running (1.188 ± 0.12) (Figure 6c) however neither were significantly different than controls (1.16 ± 0.05).

When assessing lymphatic clearance, we observed a significant reduction (p = 0.0003 and p = 0.01) in the initial change in fluorescent intensity in running post-injection (−0.19 ± 0.05) compared to pre-injection (0.36 ± 0.1) and non-running controls (0.20 ± 0.1) (Figure 6b). however, we did not observe any significant change in tau for lymphatic tracers for the various conditions of running (Figure 6d). These data suggest the transient effect of running may be lost over the much longer timescale for which lymphatic clearance occurs.

## Discussion

In this manuscript, we demonstrated the ability of multichromatic NIR imaging’s ability to assess interstitial clearance mechanisms from multiple tissue beds. Clearance pathways and rates are essential in tissue homeostasis and dictate how biomolecules interact with their intended targets. Clearance to lymphatics or venous circulation has is understudied *in vivo*. Using tissue phantoms, we established the exposure time and tissue depth limitations required for *in vivo* imaging. We determined that sensitivity/cross talk between the dyes did not exist, confirming that the changes in signal intensity resulted from changes in concentration or position of our tracers within our ROIs. We then used the mouse tail, a tissue drainage bed with well-defined physiology, to show that we could target lymphatic and venous circulation and quantify function. Lastly, we demonstrated the capacity to quantitatively image routes of clearance from the joint space. We utilized an exercise-based intervention to confirm that the technique had the sensitivity to assess clearance changes.

Our study demonstrated the size dependence of interstitial molecules via venous and lymphatic pathways via simultaneous imaging. Proulx et al. showed that NIR tracers that clear through lymphatics have a delayed uptake into the systemic circulation compared to molecules that drain directly into the blood stream^18^. In our moues tail study, the free dye intensity in the blood ROI is initially low. That signal intensifies in the tail vein over time, likely due to a renal clearance not surpassing the intradermal depot clearance over this total imaging window^32^. Thus, this dye’s intensity in the blood circulation continuously increases as the concentration delivered to the blood over time increases. The PEG tracer signal intensity traces exhibited the characteristic phasic contractions attributed to lymphatic pumping. Therefore, our mouse tail experiment validated our two tracers’ size-based partitioning to distinct routes of clearance.

The joint space is a unique interstitial space comprised of synovial fluid—hyaluronic acid, lubricin, and filtered serum—that hydrates the joint tissues and buffers the outflow of materials from the joint space^33,34^. A solute that leaves the joint space must diffuse through the synovial fluid, then into the synovial membrane. The synovial membrane is a specialized tissue that retains the synovial fluid while also housing the venous and lymphatic fluid exchange machinery to clear solute from the joint^25,35^. Smaller materials can more easily diffuse through the synovial fluid matrix and thus exit the joint space faster^36^. Larger molecules can more easily entangle in the synovial fluid matrix and therefore have longer residence times within the joint space^37^. Using multichromatic imaging with sized tracers enables quantifying venous and lymphatic clearance kinetics in the joint, simultaneously rather than separately as done in the previous studies^38^ and furthers the ability to determine the relationship between lymphatic and venous uptake *in vivo*.

Exercise has been shown in previous studies to increase interstitial^39^, venous^40^, and lymphatic^41^ flow to the muscle. In this study, we used exercise as an intervention to demonstrate the sensitivity of multichromatic NIR imaging to measure changes to venous and lymphatic clearance. In the joint space, exercise and joint loading have increased intra-articular pressure and cartilage flux. In this study, we showed that injection that was followed by exercise transiently increased lymphatic outflow from the joint; however, exercise did not have a significant effect on venous clearance. Interestingly, running pre-injection led to delayed clearance of both free dye and PEG, as exhibited by the presence of a larger peak intensity from the joint than their respective controls, which could be a consequence of altered hydrodynamic forces or delayed dye dispersion.

Our current setup limited our temporal sampling frequency in both in vivo experiments. For running experiments, sampling frequency was limited by the time required for an animal to recover from and back under anesthesia. An ideal setup would be a wearable sensor that would go around the knee, which allows us to see the concentration in real-time without anesthesia. Similarly, in capturing the routes of clearance in the tail, we could only use one channel at a time due to our stereoscope’s filter imaging limitations. Two cameras and light paths, or a computerized filter wheel, would simultaneously assess these two tracers with higher temporal frequency to quantify the relationship between vascular and lymphatic uptake *in vivo*. Additionally, the tracer sizes were designed to evaluate the particulate transport within fluid; however, there are also cell-mediated mechanisms by which transport occurs *in vivo*, which could be imaged using these multichromatic approaches.

We conclude that multichromatic NIR imaging is capable of simultaneous imaging of lymphatic and venous-mediated fluid clearance with great sensitivity and can be used to measure transient changes in clearance rates and pathways. The fluorophores and materials could be refined to provide more colors and construct sizes in the NIR range for 3 or 4 color imaging. The NIR-II imaging window could be used to visualize deeper structures *in vivo*. This methodology can be applied in future studies that assess the effects of diseases or surgical interventions on interstitial solute transport and tissue fluid homeostasis.

---

**Supplementary Video 1:** Lymphatic uptake of PEG in mouse tail in four discontinuous imaging windows post-intradermal injection.

**Supplementary Video 2:** Venous uptake/concentration of free dye over four discontinuous imaging windows post intradermal injection.

**Supplementary Video 3:** Superimposed videos of lymphatic and venous uptake of PEG and free dye in the mouse tail.

## Data and materials availability

The raw data for this study were generated at Georgia Tech and Emory University. Data or materials supporting this study’s findings are available from the corresponding authors BD and NW.

## Acknowledgments

We are grateful to Dr. Elefteria Michalaki, who carefully reviewed this manuscript. The veterinary staff at the Veterans Affairs Medical Center Veterinary Medical Unit and the Georgia Tech Physiological Research Lab for assistance. This research was funded in part by funding from the Department of Defense PRMRP grant PR171379 and the National Institutes of Health National Heart, Lung, and Blood Institute grant R01HL133216-02.

